# Buforin II-*Escherichia coli’s* DNA interactome: Detailed biophysical characterization revealed nanoscale complexes likely formed by DNA supercoiling

**DOI:** 10.1101/2022.05.20.492836

**Authors:** Daniela Rubio-Olaya, Javier Cifuentes, Paola Ruiz-Puentes, Octavio A. Castañeda, Luis H. Reyes, Jorge Duitama, Carolina Muñoz, Juan C. Cruz

**Affiliations:** Department of Biomedical Engineering, Universidad de los Andes, Bogotá, Colombia; MicroCore Microscopy Center, Vice-Rectory for Research and Creation, Universidad de los Andes, Bogotá, Colombia; Grupo de Diseño de Productos y Procesos, Department of Chemical and Food Engineering, Universidad de los Andes, Bogotá, Colombia; Department of Systems and Computing Engineering, Universidad de los Andes, Bogotá, Colombia

**Author notes:** Corresponding autor (JCC).

**Keywords:** Buforin II, peptide, interactome, nanobioconjugate, *Escherichia coli*

## Abstract

Antimicrobial peptides (AMPs) have emerged as exciting alternatives to the alarming increase of multiresistant bacteria due to their high activity against them through mechanisms that are thought to largely avoid resistance in the long term. Buforin II (BUFII) is an antibacterial peptide hypothesized to kill bacteria by crossing their membranes to interact with intracellular molecules and interrupt key processes for survival. In particular, interactions with DNA have been considered crucial for triggering cell death mechanisms. However, such interactions are still unknown, and thus far, no reports are available describing BUFII-DNA complexes. Here, we describe a complete biophysical study of the interaction between BUFII and *Escherichia coli* gDNA via spectrofluorimetric, spectroscopic, and microscopic techniques, complemented with whole-genome sequencing. The *E. coli*’s DNA-BUFII interactome was isolated by an *in vitro* pull-down method aided by BUFII-magnetite nanobioconjugates. Our results demonstrated that DNA-BUFII formed round-shape nanoscale complexes by strong electrostatic interactions, likely occurring nonspecifically throughout the entire bacterial genome. Further sequencing of the isolated DNA fragments corroborated this notion and led to hypothesize that BUFII is possibly responsible for inducing DNA’s supercoiling.

Other evidence for this idea was provided by the significant DNA conformational changes observed upon interaction with BUFII. Even though the evidence found fails to describe the complete action mechanism of BUFII *in vivo*, our findings pave the way to engineer DNA-peptide supramolecular complexes very precisely, which might find application in the field of gene therapy delivery.

## Introduction

The development of new treatments for infections caused by multidrug-resistant bacteria has been a challenge for decades due to the remarkable adaptive capabilities of bacteria. The broad spectrum and bactericidal activity of antimicrobial peptides (AMPs) make them promising candidates for treatments against infection (1).

Buforin II (BUFII) is a histone-derived peptide found in the stomach of the Asian toad *Bufo gargarizans* and the skin of several South American frogs (2). BUFII has 21 amino acids forming helical structures at the N- and C-terminals. The C-terminal region has been thought to contribute to BUFII’s antimicrobial activity, giving the peptide a stable amphipathic helical structure for forming supramolecular peptide-lipid complexes. These complexes eventually lead to transient pore formation and membrane penetration, subsequently binding to DNA and RNA, blocking vital functions, and causing cell death (3). This strong interaction tendency has been attributed to BUFII’s positively charged arginine and lysine residues (4, 5). Several studies have focused on the ability of BUFII to control infections by introducing small mutations in its structure, and it has been found that it exhibits interaction with bacterial DNA (6-10). However, the knowledge about the type of interactions predominating between BUFII and DNA remains largely unknown, and whether such interactions correspond to specific binding to motifs associated with bacteria’s vital processes is undetermined.

To improve BUFII’s properties in physiological environments, nanobioconjugates of BUFII and magnetite (Mgnt) have been previously developed, where BUFII was immobilized on Mgnt nanoparticles to enhance its stability without significantly altering its membrane-translocating and antibacterial properties. Also, these strategies have enabled drug delivery and medical images associated with the use of magnetic fields that exploit Mgnt’s responsiveness to such stimuli (11, 12). These nanobioconjugates are also a promising innovation tool to enable pull-down molecular techniques because of their unique interactions with proteins or peptides *in vitro* (13). Additionally, physicochemical methods previously used in translocating peptides analysis are valuable for understanding the mechanism of action of AMPs, including advanced spectroscopic and microscopic techniques for analyzing the complexes formed by peptide-DNA interactions, determining the underlying structure-activity relationships, and monitoring secondary and tertiary structural changes (14-16).

This study analyzes the interaction between the peptide BUFII and genomic DNA of *Escherichia coli* via molecular, spectroscopic, and microscopic techniques complemented with whole-genome sequencing. We showed the ability of BUFII-Mgnt nanobioconjugates to enable pull-down assays to enrich and isolate the interactome of interest. Sequencing data validated the use of BUFII-Mgnt to isolate the interactome and provided insights into the types of interactions with gDNA *ex vivo* that are mainly unspecific. This information was supported by microscopic evidence showing the assembly of nanoscale supramolecular structures as DNA and BUFII complexed together, which can be most likely related to the intricacies of random interactions. We found significant conformational changes in the secondary structure of BUFII upon contact with DNA and demonstrated that electrostatic forces are mainly responsible for such strong BUFII-DNA interactions. Taken together, our results led us to hypothesize that DNA supercoils, through interactions with BUFII, can be further exploited for engineering novel supramolecular complexes for applications in gene delivery.

## Materials and methods

### Extracting *E. coli* genomic DNA

Genomic DNA from *E. coli* K12 (F-lambda-*ilvG*-*rfb*-50 *rph*-1) was extracted by ultrasonication according to the protocol described previously by Zhang et al. (17). Bacteria were cultivated in LB media overnight and centrifuged to discard the medium. Then, cells were washed with TE buffer (10 mM tris HCl, 1 mM EDTA, pH 8), and 1.5 mL of sonication buffer (50 mM tris HCl, 10 mM EDTA, 100 ng mL^-1^ RNAse, pH 7.5) was added to the pellet along with 1 g of glass beads in 2 mL Eppendorf® tubes. The samples were vortexed for 5 minutes to disrupt cellular membranes and release the genomic content. After centrifugation, the supernatant was transferred to 0.5 mL Eppendorf® tubes and sonicated in cold water into an ultrasonic bath (BRANSONIC 5510R-DTH, Danbury, CT, USA) in 1-minute cycles, six times. Samples were then incubated for 30 minutes at 80°C to denature proteins and centrifuged to collect supernatant with DNA fragments of approximately 500 bp in size. To evaluate the purity of the extracted DNA, it was assessed on the optical density ratio of 260 and 280 nm (OD260/OD280 = 1.8). Additionally, DNA concentration was determined by measuring the absorbance at 260 nm in a spectrophotometer (Denovix Inc, New Castle County, Delaware, USA) at room temperature.

### Sample preparation

Buforin II-Magnetite (BUFII-Mgnt) nanobioconjugates were synthesized as previously described (11, 12) to form an aqueous suspension with a 1.6 mg mL^-1^ final concentration, containing approximately 32 mg mL^-1^ of BUFII. The complexes were prepared by mixing DNA and nanobioconjugates to obtain different ratios between anionic charges of the *E. coli* K12 DNA fragments and cationic charges of the peptide (N+:P− ratio). Complexes prepared with the nanobioconjugates and DNA were incubated at 4 °C for 1 hour under constant agitation.

Additionally, complexes of BUFII and DNA were prepared by adding a binding buffer (5% glycerol, 10 mM tris HCl, 1 mM EDTA, 1 mM DTT) to the mixture, followed by incubation at 4 °C for 30 minutes under agitation.

### Agarose gel electrophoresis

Agarose gel electrophoresis was carried out to verify DNA extraction and BUFII interaction with DNA. Samples were solubilized in 6X loading buffer (New England Biolabs Inc., Ipswich, USA) and then separated by migration on a 1% agarose gel dissolved in TBE buffer (10.8 g L^-1^ tris, 0.5M EDTA, 5.5 g L^-1^ boric acid, pH 8) in the presence of HydraGreen (3 mL, ACTGene, USA) as the intercalating agent. Electrophoresis was carried out for 1 hour at 90 V using an electrophoresis system (mini–DNA SUB CELL, Bio-Rad, Mississauga, Canada). The DNA fragments were visualized in a Gel Doc XR+ Gel Documentation System (Bio-Rad, Mississauga, Canada) and photographed.

### Pull-down and next-generation sequencing (NGS)

To obtain the *E. coli* K12 gDNA fragments that interacted with BUFII, complexes with the nanobioconjugates were prepared as described above. As shown in Fig 1A, particles were precipitated with a magnet, and the supernatant was removed. The complexes were washed with sterile distilled deionized water twice to remove non-interacting fragments. BUFII/DNA complexes were then eluted by adding 100 μL of elution buffer (1% SDS, 0.1 M NaHCO3, pH 8), vortexed for 10 minutes, and spun down at 13,000 RPM for 5 minutes. A negative control assay was performed using Mgnt nanoparticles without BUFII instead of nanobioconjugates to dismiss unspecific interactions. The supernatant with 100-500 bp DNA fragments was purified with Monarch PCR & DNA Cleanup Kit (New England Bioloabs Inc., Ipswich, USA) and stored at -80 °C until further use. Whole-genome sequencing was performed on the Illumina NovaSeq platform using Truseq ChIP-seq library and protocols (Illumina, San Diego, CA, USA) with a minimum Phred quality score of 30. A total production of 2 Gbp on raw data for each sample was obtained from the sequencing process, which corresponds to a genome-wide average depth of 400X. Two samples were sequenced, a sample obtained from the previously described interaction assay and a negative control performed as described previously, but in the absence of BUFII to detect the nonspecific interactions between DNA and bare Mgnt.

**Fig 1.**
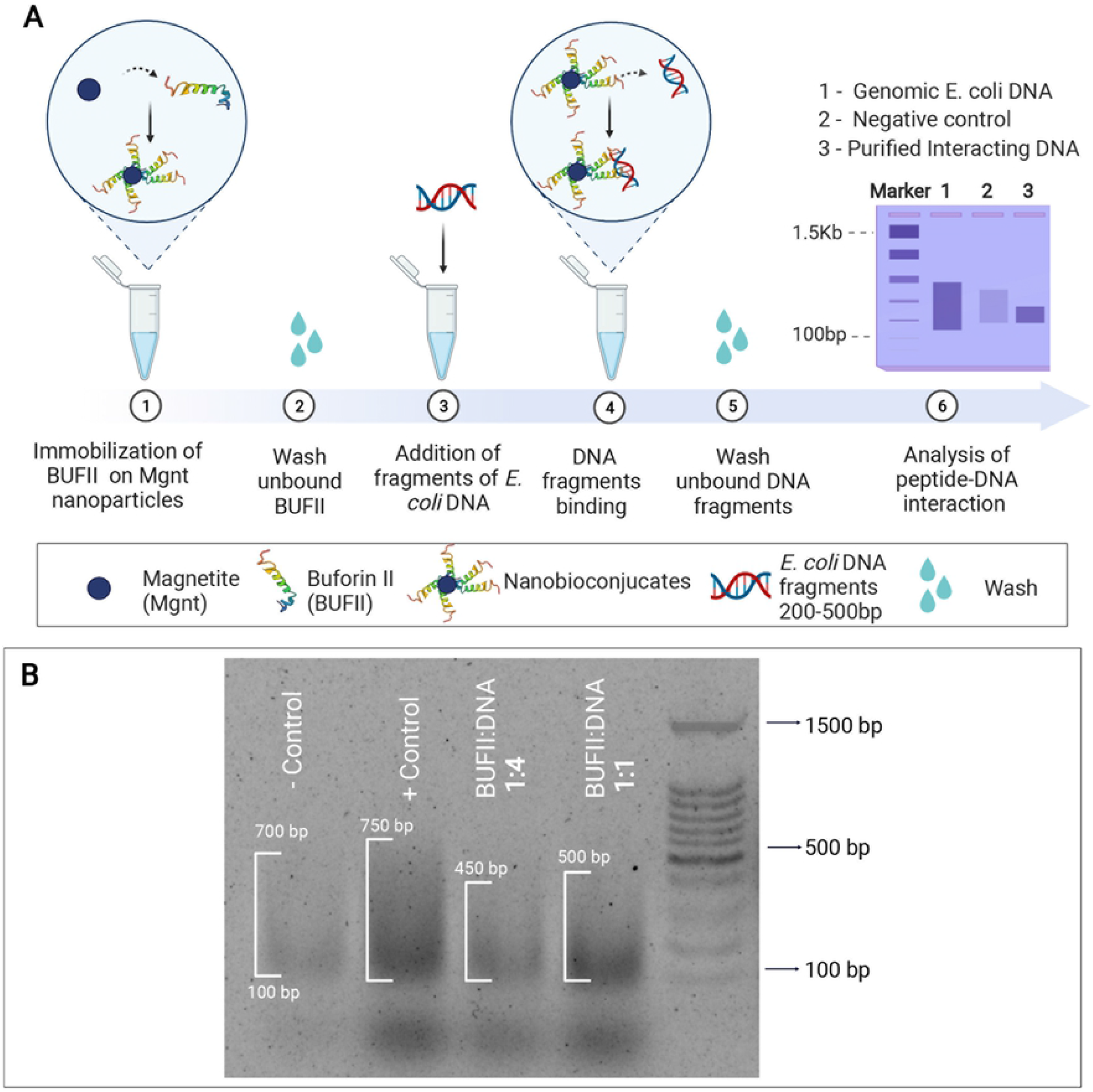
BUFII interactome pull-down description and validation. (a) Schematic of BUFII-DNA pull-down assay using magnetic properties of magnetite (Mgnt) nanoparticles in BUFII-Mgnt nanoconjugates to isolate interacting fragments of *E. coli* DNA. Created with BioRender.com (b) Agarose gel for pull-down verification. BUFII:DNA1:4/1:1, isolated fragments from interaction with Mngt-BUFII nanobioconjugates with DNA fragments. (-) Control, isolated fragments from interaction with magnetite nanoparticles. (+) Control, genomic DNA fragments.

### Analysis of sequenced data

The genome assembly of the *E. coli K12* strain with accession ID AP009048.1 was downloaded from the NCBI Nucleotide database and used as a reference for the analysis. Raw reads were mapped to the genome using bwa mem v. 0.7.12 with default parameters (18). Windows of 100 bp showing significant differences in reading depth between the treated and the control sample were identified running the ReadDepthComparator command of NGSEP v4.1.0 (19). Windows with a p-value below 10^−6^ and log fold-change above 4 were selected. Windows separated by less than 1 kbp were merged to identify the peaks of differential read depth. Bedtools v2.30.0 was used to generate FASTA files from the coordinates in pair bases corresponding to each peak. Results were visualized in the Integrative Genomics Viewer. Sequences from the FASTA files were searched using the BLAST tool from NCBI (20) to identify encoding genes in bacteria and provide insights into the possible biological functions altered by the interaction with BUFII.

### Fluorescence measurement

The fluorescence spectra of *E. coli K12* genomic DNA in the presence and absence of BUFII were measured using a FluoroMax4 spectrofluorimeter from HORIBA Scientific at room temperature as described previously (21). Data were collected using the FluorEssence® software. The reference excitation monochromator and detector were previously verified and calibrated with water and rhodamine B to avoid temporal fluctuations in the source during excitation scans. The excitation wavelength was set at 535 nm, and emission spectra were measured between 550 and 750 nm for increasing BUFII concentrations (0, 9, 18, 36 μg mL^-1^) to a DNA fixed concentration (25 μg mL^-1^). The measurements were conducted for both BUFII-Mgnt/DNA complexes and BUFII/DNA complexes.

#### Competitive binding of BUFII and SybrGreen (SG) with bacterial DNA

Additional fluorescence measurements were carried out to analyze competitive binding. Assays were carried out in denatured water containing a fixed concentration of SybrGreen (3 μg mL^-1^)-DNA (25 μg mL^-1^) with varying concentrations of BUFII (0, 9, 18, and 32 μg mL^-1^). The SG–DNA solution was incubated at room temperature for 10 min to allow the mixture to reach equilibrium. Then, BUFII-Mgnt/BUFII were added to the SG–DNA solution, and the fluorescence spectra were recorded between 500 and 700 nm for each test solution with an excitation at 490 nm.

The competitive binding of SG against BUFII and bacterial DNA was also probed via fluorescence measurements. The assays were carried out in denatured water in the presence of a fixed concentration of BUFII (18 μg mL^-1^)–DNA (25 μg mL^-1^) and varying the concentrations of SG, making serial dilutions (1:2) of the previous concentration (0, 0.75, 1.5 and 3 μg mL^-1^). The BUFII/BUFII-Mgnt-DNA solution was incubated at room temperature for 10 min to reach equilibrium. Then the solution containing SG was added to the BUFII/BUFII-Mgnt–DNA solution. The solutions were excited at 490 nm, and the emission spectra were recorded from 500 to 700 nm.

### Fourier-Transform Infrared Spectroscopy (FTIR)

Samples were prepared as described previously. BUFII-Mgnt nanobioconjugates were precipitated to discard the supernatant and dilute the complexes in deuterated water. Samples were examined on a Bruker Alpha II FTIR Eco-ATR instrument (Bruker Optik GmbH, Ettlingen, Germany). The peptide concentration was kept at 32 μg mL^-1^, and the corresponding amount of *E. coli K12* DNA fragments was added to match the desired molar ratios. Droplets from solutions were placed on top of the instrument’s germanium crystal, and data were collected between 4000-600 cm^-1^ with a spectral resolution of 2 cm^-1^. After measurements, FTIR curves were smoothed out for noise suppression by replacing the intensity value of each data point with the value obtained by averaging the intensities of three points. The measurements were performed for both BUFII-Mgnt/DNA complexes and BUFII/DNA complexes.

To analyze changes in the secondary structure of BUFII upon complexing with DNA, the Amide I and II (1800 cm^-1^ -1500 cm^-1^) spectra were deconvoluted aided by the second derivative of the spectra as described previously by Kong and Shaoning (22).

### Transmission Electron Microscopy (TEM)

TEM imaging (FEI, Tecnai F20 Super Twin TMP, Hillsboro, OR, USA) was carried out for complexes with both immobilized and free BUFII and *E. Coli K12* DNA fragments (1:1). The samples were fixed onto lacey Carbon grids by depositing drops of the main solution (0.1 mg mL^-1^) on the top and letting them rest for 5 minutes. Next, uranyl acetate staining was performed by adding 5 μL of 2% uranyl acetate solution and leaving the sample to rest for 5 minutes at room temperature. The microscope was operated at a 200 keV acceleration voltage to visualize the samples.

### Atomic Force Microscopy (AFM)

Topography AFM images were obtained using an MFP3D-BIO AFM (Asylum Research, Santa Barbara, CA, USA) instrument. As described by Pillers et al., samples were prepared to immobilize complexes with both free and immobilized in Mgnt BUFII and *E. coli K12* DNA fragments (1:1) on silicon substrates. Silicon surface chips were previously cleaned using RCA1 (28 % ammonium hydroxide, 30% hydrogen peroxide, DI water) and RCA2 (14% hydrochloric acid, 30% hydrogen peroxide, DI water) solutions (23). Silicon chips were functionalized with a 2% (3-aminopropyl) triethoxysilane (APTES, 98%, Sigma-Aldrich, St. Louis, MO, USA) solution to attach complexes to the surface by interactions with free amine groups. The main sample solution was then mixed, and 4 μL were deposited on the silicon substrate. Samples were left to rest for about 10 minutes and then rinsed with 100 μL of sterile 18 MΩ x cm water and dried with N_2_ for 1 min. The samples were stored in a clean container until further experiments to avoid possible artifacts during imaging. All measurements were conducted with an AC240TS cantilever (Oxford Instruments, Asylum Research, Santa Barbara, CA, USA) using the tapping mode set at a 70% free amplitude setpoint. The images were obtained using different scan sizes in a range from 1.5μm x 1.5μm up to 5μm x 5μm at a resolution of 1024 by 1024 pixels and with a scan rate of 1 Hz.

## Results and discussion

### Nanobioconjugates allowed obtaining the BUFII interactome with DNA

Pull-down experiments aided by BUFII-Mgnt nanobioconjugates allowed the successful isolation of *E. coli* K12 DNA fragments that interacted with immobilized BUFII, as described in Fig 1A. The interactome showed in the electrophoresis gel (Fig 1B) corresponds to assays performed with DNA fragments ranging from 200 bp to 1 kbp. However, it was possible to isolate only those between 100 and 500 bp. Also, it was found that Mgnt shows some level of interaction with DNA, which is most likely due to electrostatic interactions with charged functional groups present on the surface of Mgnt. We performed direct sequencing of the DNA present in the experiment, as well as the DNA in the control experiment. As shown in Fig 2A, data from sequencing were successfully mapped to the K12 reference genome and visualized to determine whether the proposed method is sufficiently robust to isolate the interacting *E. coli K12* DNA fragments accurately. The panel shows an example of the depth distribution around the bcsA gene. Although the depth is around the expected average of 400X, important differences are observed between samples, indicating that BUFII interaction with DNA is distinguishable from that between DNA and bare Mgnt. Additionally, regions of significantly higher read depth were identified for the library of the BUFII-containing sample, which in comparison with the control, suggest the presence of specific interactions.

**Fig 2.**
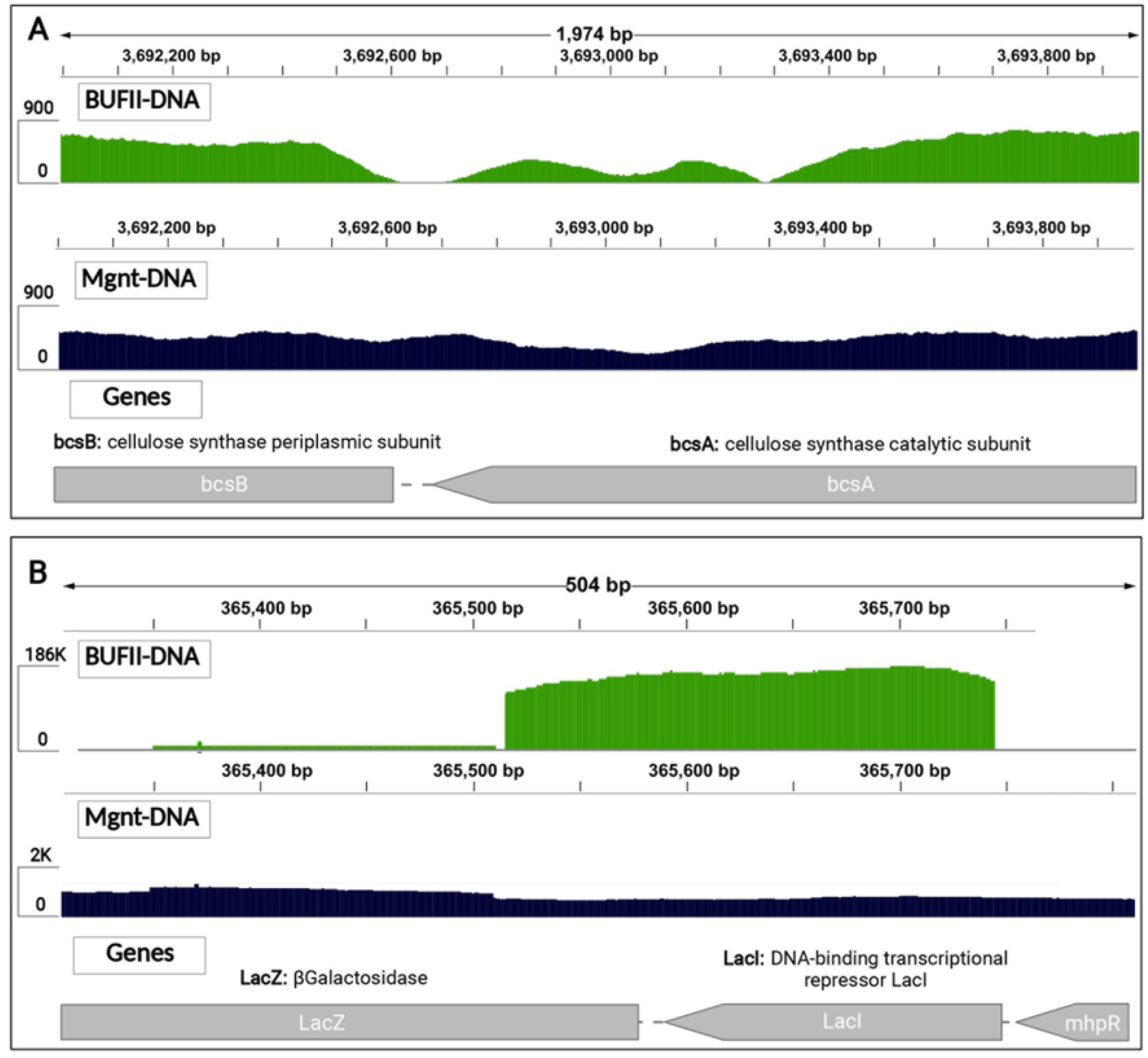
Sequencing data from BUFII interactome visualization. Data from interactomes was mapped and visualized to identify motifs associated with the antimicrobial activity of BUFII using interaction between *E. coli K12* DNA and Mgnt as a control. (a) Visualization of a fragment of mapped reads showing differences in the enrichment patterns between samples. (b) Sample from interaction with BUFII showed important enrichment in the genome coordinates associated with LacI repressor gene compared to Mgnt sample.

### BUFII-DNA interaction is mainly unspecific but appears to have an affinity towards some DNA sequences

In general, the BUFII containing sample showed regions over the whole genome with significantly high depth (above 1000X). Conversely, no coverage was obtained in some other regions, an observation that will be studied further in our future contributions. Fig 2B shows the most interest difference in read depth between samples. The BUFII containing sample has a zone of enrichment that reaches approximately 130000X, indicating a particular affinity of the peptide towards the indicated 400 bp region, which was not observed in the Mgnt control.

We then proceeded to search for peaks indicating a higher affinity of the peptide for specific regions of the genome. The most relevant ones correspond to the same zone of higher enrichment between 365401 and 365800 bp. The sequences in FASTA format were recovered from the peak positions to evaluate their biological function and whether it was related to the antimicrobial activity of BUFII. The analysis of the sequences shows that this region encodes for the Lac operon and specifically for the LacI repressor and the β-galactosidase enzyme. The function of this region is to switch the metabolic pathway from glucose to lactose when the bacterium is in a medium with low glucose content but rich in lactose or any of its derivatives (24).

Additionally, the MEME suite software version 5.4.1. (25) was used to try to identify motifs enriched at the regions with differential read depth. Unfortunately, the number of regions with significant differences in read depth was insufficient to predict with enough statistical significance specific interactions with DNA motifs. However, this initial interaction experiment suggests that despite the largely unspecific interaction between BUFII and phosphate groups in the DNA backbone chain as has been described previously (7, 9), the peptide showed a surprisingly greater affinity towards some specific DNA sequences. Our data is not conclusive to explain this result. Hence, we are planning to conduct further in silico and experimental studies that involve MD simulations and calorimetric measurements of the interacting complexes.

To validate the data related to the enriched zone corresponding to LacI repressor in the *E. coli* K12 Lac Operon, an assay described in the supplementary material for inducing GFP expression with the BUFII-Mgnt nanobioconjugates was performed. The obtained results demonstrated that the BUFII-Mgnt inhibited the LacI repressor, leading to GFP expression (S1 Fig). Despite confirming the particular affinity of BUFII towards this sequence, the experiment failed to explain the potent antimicrobial activity of BUFII because there is no evidence of metabolic alterations that compromise vital bacterial functions. Sequencing data and the validation experiment let us elucidate that BUFII might have an affinity for some DNA motifs; however, the underlying interaction is mainly unspecific.

### BUFII-DNA interaction is mainly mediated by electrostatic interactions

#### BUFII-DNA interaction

Fluorometric assays performed with complexes containing BUFII and gDNA fragments isolated from *E. coli K12* (0–32 μg mL^-1^) provided additional insights into the possible underlying mechanisms for the observed interactions.

As shown in Fig 3A, in the absence of BUFII, the DNA of *E. coli K12* emitted a characteristic fluorescence profile in water at room temperature, with a peak close to 585 nm after excitation at 535 nm. Upon adding BUFII-Mgnt into the DNA solution, the fluorescence intensity decreased dramatically, most probably due to the ability of Mgnt nanoparticles to absorb the fluorescence emitted by the BUFII-DNA complexes or by the steric hindrance caused by the Mngt nanoparticles that prevent sufficient DNA and BUFII encounters for a sustained interaction that can be monitored spectrofluorimetrically.

**Fig 3.**
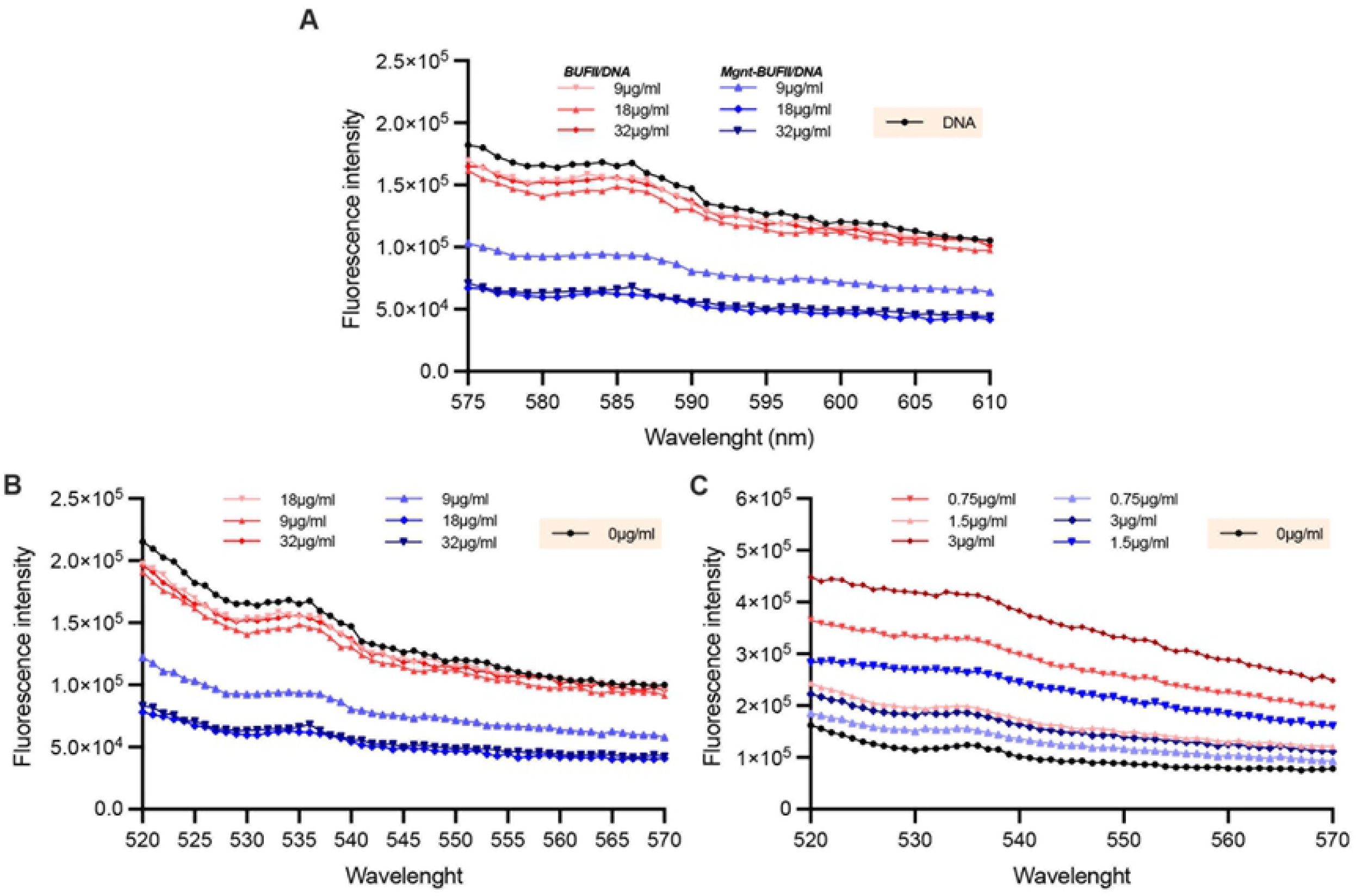
BUFII-DNA complex spectrofluorometric assays. (a)Fluorescence spectra of DNA fragments in the presence of increasing amounts of BUFII. A fixed concentration of DNA was mixed with increasing amounts of BUFII/Mgnt-BUFII. Samples were excited at 535nm. (b) Fluorescence spectra of DNA fragments and BUFII in the presence of SybrGreen (SB), a fixed concentration of DNA and SB was mixed with increasing amounts of BUFII/Mgnt-BUFII. (c) Fixed concentration of DNA and BUFII/Mgnt-BUFII was mixed with increasing amounts of SB. Samples were excited at 490 nm.

The behavior of fluorescence signals changed as the concentrations of BUFII increased. This might be primarily due to changes in the DNA backbone structure, similar to previously reported DNA clustering that reduces fluorescence intensity and decreases the collisional frequency of solvent molecules with DNA (26). Spectrofluorimetric techniques have been successfully used to detect specific DNA–protein binding because they allow direct measurement of binding in solution.

Changes in the fluorescence emission spectrum of a protein upon binding to DNA can often be used to determine the stoichiometry of binding and to decipher the types of molecular interactions (27-29). In this case, data indicate that BUFII could interact with the backbone DNA or locate deep into the DNA hydrophobic regions, evidenced by fluorescence intensity changes because of the difference in the amount of light absorbed at excitation and emission wavelengths. The results point toward an incomplete inhibition of the DNA fluorescence spectra.

#### Competitive binding of BUFII and SG with bacterial DNA

To further decipher the mechanistic details of the interactions between BUFII with DNA, a competitive binding experiment was carried out using SybrGreen (SG) as a double-stranded DNA (dsDNA) intercalating agent (λ_ex_ = 490 nm and λ_em_ = 500–700 nm). This method was used to assess the ability of the peptide to prevent the intercalation of SG within DNA strands. In general, when small molecules like SG bind to dsDNA, interaction cause changes in the fluorescence spectra compared to what is observed for solutions in the absence of this ligand. However, fluorescence quenching will be observed when a second ligand competes for the DNA-binding sites (30).

As shown in Fig 3B, the addition of BUFII to dsDNA pretreated with SG caused a noticeable fluorescence quenching, indicating that BUFII competed with SG in binding to DNA. This observation suggested that BUFII replaced some SG molecules that interacted with the DNA base pairs and released them into the aqueous solution. Consequently, a decrease in the emission was observed (31). However, fluorescence spectra for each sample show that the reduction in fluorescence is not directly proportional to the concentration of BUFII/BUFII-Mgnt. At a concentration of 18 mg mL^-1^, the fluorescence signal was reduced, suggesting that the interaction between the molecules is most likely mediated by electrostatic forces, which, after reaching an equilibrium (cation:anion), might favor the ability of the peptide to intercalate between adjacent base pairs and replace the SB molecules (32).

Further support for the proposed mechanism of BUFII binding to DNA via intercalation was given through a competitive binding assay of SG with bacterial dsDNA against BUFII (Fig 3C). With the addition of SG, the characteristic fluorescence band of the BUFII–DNA complex with a maximum at about 535 nm (excited at 490 nm) rose gradually, indicating that some of the SG molecules intercalated into the DNA base pairs instead of BUFII. SG–DNA replaced BUFII–DNA gradually, showing that, in this case, an increase in fluorescence intensity was proportional to SG concentration. This supports previous information on the specific interaction between peptides and bacterial DNA, pointing towards electrostatic interactions mediated by the existing charge differences.

To dismiss the effect of Mgnt nanoparticles in the interaction between DNA and BUFII, negative controls for all Mgnt-BUFII nanobioconjugates concentrations were performed by adding bare Mgnt (S2 Fig). The obtained data suggest an interaction between Mngt nanoparticles and DNA, as evidenced in the pull-down assay; however, the observed changes in the fluorescence spectra of DNA are negligible in the presence of Mgnt. Additionally, information obtained for BUFII-Mgnt nanobioconjugates was consistent with that of BUFII alone, confirming that even after immobilization on nanoparticles, BUFII can still interact with bacterial dsDNA very strongly.

### Changes in BUFII-DNA complexes’ secondary structure

FTIR assays were carried out to identify changes in the secondary structure of both DNA and BUFII when forming complexes and to gain insights into the chemical groups playing a role in complexation. The vibrational spectra of the samples showed that specific functional groups appear responsible for the association between BUFII and DNA. Also, they reveal important information on the impact of charge changes in complexation for each ratio (+:-) studied.

Experiments were focused on two major regions of the infrared spectrum, the sugar− phosphate region of DNA and the amide I and amide II bands of the peptide. The first range encompasses wavenumbers between 950 and 1200 cm^-1^, which carries information on chemical bonds associated with the DNA phosphate and ribose groups. In contrast, the second region locates between 1500 and 1800 cm^-1^ and encompasses vibrations related to NH moieties on peptide backbones. Also, vibrations associated with DNA guanine groups can be found between 1700 and 1800 cm^-1^.

#### Sugar-phosphate region

Data from the sugar− phosphate region shown in Figs 4A and 4C reveals the effect of complexation on the DNA backbone. There are vibrations related to involved functional groups as a function of the evaluated charge ratios. Particularly, bands associated with stretching of C− O bonds of deoxyribose and stretching of − PO2− groups located at 1033 and 1084 cm^-1^ (33). Conformational changes in DNA’s backbone are evidenced by peak shifts between 1084 and 1048 cm^-1^, indicative of interactions between DNA phosphate groups and BUFII. Changes in the 1084 cm^-1^ peak are related to the reorganization of groups adjacent to phosphates, indicating considerable conformational changes in the DNA strands (34).

**Fig 4.**
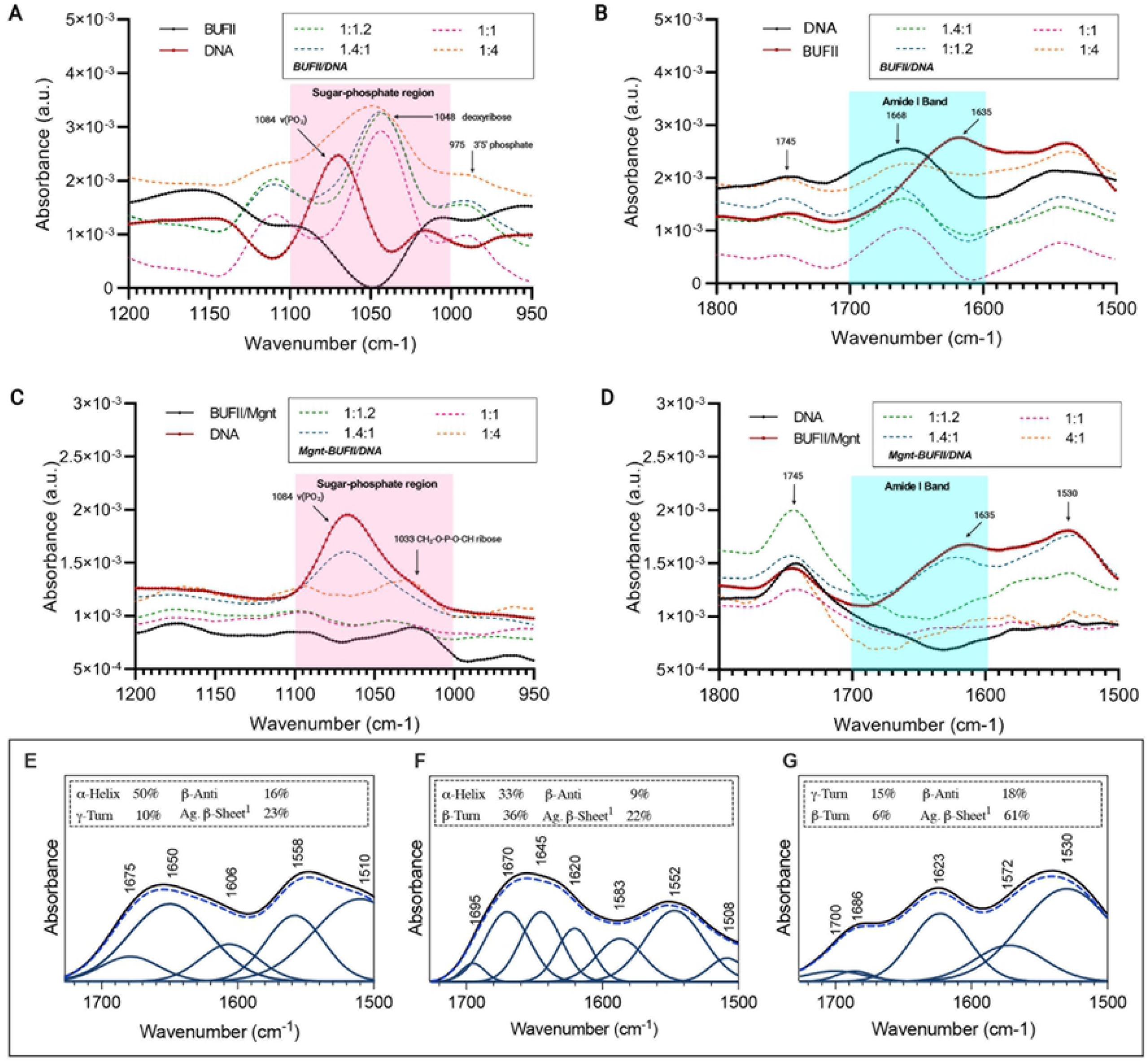
Spectroscopy assays from solutions containing BUFII/Mgnt-BUFII and genomic *E. coli K12* DNA at different molar ratios. (a, c) FTIR data collected across spectral range corresponding to the sugar-phosphate region [950-1200 cm^− 1^]. (b, d) FTIR data collected across spectral range corresponding to the peptide amide I band [1800-1750 cm^− 1^]. (e, f, g) Spectra of the Amide I and carbonyl stretching region were deconvolved into component sub-bands to analyze secondary structural changes of BUFII aided by the second derivative of FTIR spectra for BUFII (e), BUFII/DNA 1:1 complex (f) and BUFII/DNA 1:4 complex (g).

When comparing the complexes’ bands with free DNA spectra, the shifting in the peak at 1048 cm^-1^ seems related to interactions occurring at charge ratios closer to neutrality. This supports the findings based on fluorescence for the peptide-DNA interaction mediated by electrostatic forces discussed above. Taken together, our data suggest that it is very likely that BUFII/DNA complex formation proceeds by polyelectrolyte brush states that result in unique 3D topologies (possibly due to different BUFII interacting conformers, see below), as has been reported previously (35).

#### Amide I and II region

Figs 4B and 4D show infrared spectra across the amide I region. In this case, solutions containing complexes closer to the neutrality exhibit spectra with peaks with increased intensity and sharpness, whereas those with the prevalence of cationic charges show a significant intensity reduction (36).

These changes may also suggest the appearance of new peptide conformers upon complexation with DNA. Differences in the characteristic amide band between 1700 and 1600 cm^-1^ are related to beta-sheet conformations and disordered structures associated with random coil and clustered conformations (37).

#### Deconvolution of Amide I and carbonyl stretching region

To identify the secondary structural changes in the formed BUFII/DNA complexes, we analyzed the amide I and carbonyl stretching region (1500-1800 cm^-1^), deconvolving each spectrum with the aid of the second derivative. Secondary structure was estimated under the principle that absorbance reflects the backbone conformation of proteins maintained by hydrogen bonding. Consequently, the deconvolved sub-bands in the amide I region can be correlated with α-helices, β-sheets, turns, and random coils (22).

BUFII spectra, when interacting with DNA, are shown in Fig 4E. The Amide I band was deconvolved into three sub-bands: the first one peaking at 1650 cm^-1^ and related to α-helices, the second one at 1606 cm^-1^ associated with disordered structures and aggregated beta-sheets that may arise when the peptides are dissolved in aqueous media. In this regard, it has been reported that BUFII tends to form aggregates and disordered structures different from its mainly known helical conformation (38). Finally, a third small band at 1675 cm^-1^ may be related to the disordered coil of BUFII. The α-helical conformations were confirmed by the Amide II peak at 1510 cm^-1^, while the disordered ones and beta-sheets were confirmed by a peak at 1558 cm^-1^ in the Amide II band.

When an equimolar gDNA concentration is added to the BUFII solution, drastic changes are observed in the deconvolution profile in Fig 4F, with the presence of four sub-bands showing a shift in the α-helices predominance towards aggregated structures, as evidenced by a percentage increase in the turns in the peaks at 1695 and 1670 cm^-1^. The changes indicate that the conformational changes of BUFII occur to improve interaction with DNA. This has been confirmed by published data that suggest that binding between the carbonyl groups of peptides and the phosphate groups of DNA are favored in the presence of β-sheets (39).

Fig 4G shows the FTIR deconvolution upon adding a fourfold molar concentration of DNA to the BUFII solution. A change in the spectrum indicates a predominance of aggregated β-sheets (1623 cm^-1^ peak). Also, a decrease in intensity and shift from 1552 to 1572 cm^-1^ within the amide II band strongly suggests changes in the N-H stretching mode of adenine, which can be correlated to specific interactions with DNA motifs and structural alterations along the backbone (40). This validates further the information obtained by the fluorescence studies discussed above, indicating that equimolar ratios favor charge-mediated interactions.

### BUFII-DNA complexes are nanoscale supramolecular structures

Transmission electron microscopy (TEM) imaging was conducted to provide insights into the complex formation between free and immobilized BUFII and *E. coli K12* DNA fragments. Images obtained allowed us to elucidate the morphology and size of each complex, and to elaborate more compelling arguments about the identified secondary structural changes presented above. According to the molar equivalence between positive and negative charges, samples were prepared from solutions containing peptides at a concentration of 0.1 mg mL^-1^. Micrographs shown in Fig 5A confirm complexation between the peptide and DNA. They reveal the formation of discrete nanostructures with sizes ranging from a few nanometers up to hundreds of nanometers. These structures suggest that molecules interact to form rounded structures, most likely due to the presence of BUFII interacting conformers capable of packaging DNA strands very tightly through electrostatic interactions with the phosphate groups (i.e., the phosphodioxy group), as demonstrated by the collected spectroscopic data (see above). Additionally, these interactions seem to lead to regions where DNA strands appear highly supercoiled; however, the underlying mechanisms are yet to be described.

**Fig 5.**
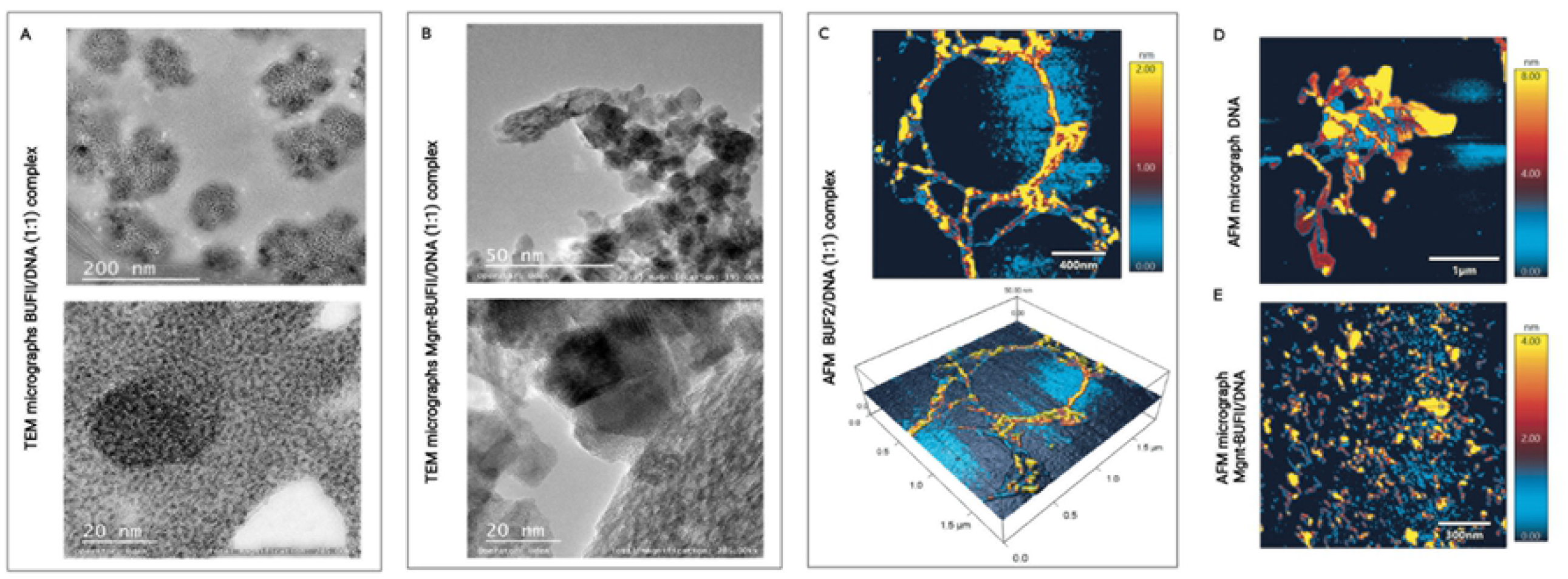
TEM and AFM micrographs from complexes formed between free and immobilized BUFII and DNA fragments at a molar charge ratio of 1:1. Presence of globular complexes of BUFII and DNA strands throughout the samples. (a) TEM micrograph for fixed BUFII/DNA complexes revealing round-shape structures. Darker spots correspond to BUFII aggregates, while DNA strands are visible as supercoiled white patches along the formed complexes. (b) TEM micrographs of complexes formed between Mgnt-BUFII nanobioconjugates and DNA. Even though no apparent structures are observable, DNA appears to coat the nanobioconjugate surfaces. (c) AFM micrograph and 3D reconstruction for BUFII/DNA complexes, round-shape structures are formed upon the interaction between molecules. (d) Shows AFM image of DNA molecule in a disordered structure by immobilization on a polished silicon wafer surface. (e) AFM image of agglomerates formed after Mgnt-BUFII nanobioconjugates and DNA interaction.

For a closer inspection of the complexes, we focused on one of the supramolecular structures formed. Fig 5A shows that most likely, BUFII locates the core of the structure, serving as a scaffold to support the DNA strands supercoiling around the aggregated peptides to give rise to rounded structures. This confirms the findings of the FTIR assays corresponding to the appearance of disordered BUFII structures and the changes in the DNA backbone associated with twisting the outer groups of the strands and, in turn, altering the planarity of the DNA molecule.

For BUFII-Mgnt nanobioconjugates, micrographs (Fig 5B) showed the nanoparticles coated by the DNA, confirming that immobilization does not affect the interacting ability of the peptide. However, it was impossible to fully identify the presence of different supramolecular complexes mainly because of the differences in the size of the analyzed structures and the nanobioconjugates.

### Imaging of BUFII-DNA confirmed strong interactions and unique supramolecular organization

To further understand the topology of the complexes, we also conducted imaging by AFM. Figs 5C and 5D show evident topological differences between the typical structure of the DNA and the formed BUFII-DNA complexes. Free DNA strands appear elongated and disordered, probably due to the immobilization protocol. After adding the peptide and immobilizing it on the silicon wafer surface at the 1:1 molar ratio, imaged complexes showed structures similar to those observed by TEM. Fig 5C revealed rounded structures of various sizes with DNA strands surrounding BUFII and forming a more extensive interconnected system.

The formed complexes show DNA strands with similar thicknesses, indicating that the peptide remains at the core while the DNA supercoils form a shell. The interaction between Mgnt-BUFII nanobioconjugates and DNA shown in Fig 5E confirms the DNA aggregation/condensation over the nanobioconjugates, as evidenced previously by the TEM imaging. However, no large structures were observable in the presence of the nanobioconjugates, suggesting that immobilization of the peptide might alter BUFII-DNA interactions, which is likely why the antimicrobial activity of BUFII reduces upon immobilization (12, 41). To our knowledge, no reports are available describing this possible DNA-peptide interaction mechanism, which suggests that this could be a new approach to studying the action mechanism of AMPs with translocating properties like BUFII.

### BUFII-DNA complexation: Structure and types of interactions

The results presented above reveal that the interaction between BUFII and DNA results in the formation of nanocomplexes of different shapes and sizes. The collected data provided several insights into the organization of molecules upon interaction and allowed us to find a rationale for the identified secondary structural changes. Moreover, we are putting forward a complete description of the assembly process for the BUFII-DNA complexes. In addition, we found compelling evidence for the significant role that the charges of involved molecules might play during this process. Finally, our findings point to a multistage assembly process occurring at different scales as larger structures appear composed of smaller interacting subunits.

Our proposal for the mechanism of complex assembly is schematically shown in Fig 6. According to this model, complexation is mainly triggered by electrostatic attraction between the negative charges of DNA phosphate groups and the cationic groups of peptide chains (Fig 6A). The initial supramolecular association is represented by the conformational changes of nucleic acid duplexes caused by interacting conformers of BUFII. The role of electrostatic forces as a significant self-assembling driving force is supported by spectrofluorimetric information in competitive DNA binding assays that suggests that BUFII could replace other strongly bound molecules. However, this type of interaction is weaker than a covalent bond, and therefore it is likely reversible (Fig 6B).

**Fig 6.**
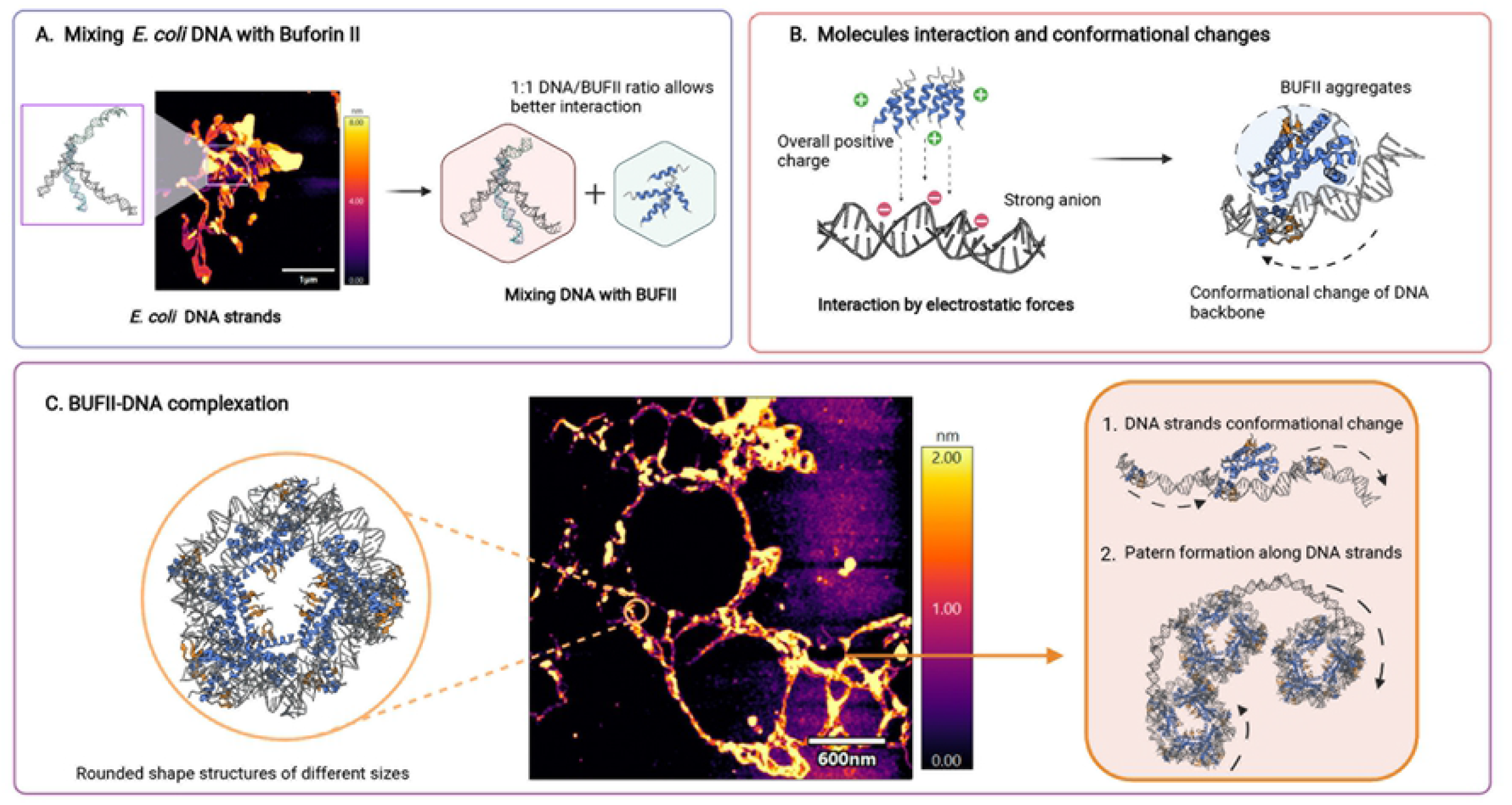
Schematic representation of the proposed interactions to form BUFII/DNA complexes. Self-assembly is most likely mediated by electrostatic forces causing conformational changes in *E. coli* DNA backbone and random agglomerated structures for BUFII. (a) Ratios that allow equimolar charges between peptide and DNA when forming the complexes result in stronger interactions. (b) Electrostatic forces appear to mediate the interactions, and secondary structural conformation changes facilitate BUFII binding to DNA. (c) Interaction between molecules results in a large complex composed of round-shape structures containing BUFII agglomerates. Strands tend to form different structures because of BUFII’s supercoiling induction.

The formation of rounded structures composed of DNA strands surrounding BUFII aggregates is illustrated in Fig 6C, along with AFM images showing large, microscopic structures in the range of 4-6 μm. Aggregates correspond to subunits found making part of larger DNA strands resulting from BUFII’s interacting conformers. The observed structures with unique topological features are hypothesized to be formed because of the supercoiling of the DNA induced by BUFII. In this regard, when DNA binds to proteins by electrostatic interactions that can constrain supercoils, negative and positive interactions lead to increased rotational dynamics and the formation of compact complexes made of the loop, thereby causing changes in transcriptional machinery (42).

Because of the presence of DNA strands in the periphery of complexes, we propose that a rearrangement of their backbone structure accompanies complex formation. This reorganization is driven mainly by interactions related to the amphiphilicity of the peptide molecules. Indeed, because BUFII contains a significant number of nonpolar groups, the resulting hydrophobic effect is likely responsible for helping the peptide locate at the cores of the supramolecular assemblies. In contrast, the high hydrophilicity of double-helix phosphates would compel DNA toward the interfaces with water (43).

## Conclusion

Previous studies of the functional mechanisms of BUFII have focused mainly on the peptide– membrane interaction, which represents the initial step of the bactericidal process. However, little is known about the BUFII–intracellular targets interaction and specifically with DNA, as it has been considered the molecular target responsible for compromising bacterial survival. Here, we reported on a novel technique to perform pull-down assays to study the BUFII-DNA interaction that takes advantage of the magnetic properties of magnetite nanoparticles, which were employed as supports for peptide immobilization. Characterization techniques showed that DNA-BUFII strong interaction leads to the formation of spherical supramolecular complexes with nanoscale dimensions. Based on the measurements of such complexes, DNA molecules appear supercoiled surrounding BUFII. Although sequencing data analysis of enriched fractions failed to provide statistically significant information regarding interaction with specific motifs that could explain the antimicrobial activity of BUFII, the enrichment of the gene for LacI binding repressor and the experimental microscopy and FTIR results provided insights into possible sequence-mediated interactions that will be explored in detail in our future contributions. Notably, the absence of reads in specific regions of the genome offers further evidence for the notion of supercoiling induced by BUFII, as such DNA structures have been reported to be difficult to sequence. Future studies will also focus on exploring the strength of peptide-DNA interactions with the aid of molecular dynamics simulations and calorimetric techniques. Our studies also provide an avenue into the rational design of peptide-DNA supramolecular structures with unique topological features, which might be applied in the engineering of novel gene delivery vehicles.

## Supporting information

**S1 Fig. Buforin II operon Lac induction validation and spectrofluorometric assay**. (a) Assay principle for the use of BUFII-Mgnt nanobiocongugates for the induction of GFP as a consecuence of the interaction between the peptide and Operon Lac genes. (b) Visualization of GFP expression in bacteria under UV light for the samples induced with different non-lethal concentrations of BUFII-Mgnt. (c) Fluorescence measurements to quantify relative GFP expression in bacteria. Experiments were conducted in triplicate to dismiss false positives.

**S2 Fig. Mgnt-DNA interaction control spectrofluorometric assays**. (a)Fluorescence spectra of DNA fragments in the presence of increasing amounts of Mgnt. A fixed concentration of DNA was mixed with increasing amounts of Mgnt. Samples were excited at 535nm. (b) Fluorescence spectra of DNA fragments and Mgnt in the presence of SybrGreen (SB). A fixed concentration of DNA and SB was mixed with increasing amounts of Mgnt. (c) Fixed concentration of DNA and Mgnt was mixed with increasing amounts of SB. Samples were excited at 490 nm.

## Notes

### Competing Interest Statement

The authors have declared no competing interest.

